# Investigating Subpopulation Dynamics in Clonal CHO-K1 Cells with Single-Cell RNA Sequencing

**DOI:** 10.1101/2024.05.22.595338

**Authors:** Luke B Morina, Haoyu Chris Cao, Alice Chen, Swetha Kumar, Kevin S McFarland, Natalia I Majewska, Michael J Betenbaugh, Winston Timp

## Abstract

Chinese Hamster Ovary (CHO) cells are used to produce monoclonal antibodies and other biotherapeutics at industrial scale. Despite their ubiquitous nature in the biopharmaceutical industry, little is known about the behaviors of individual transfected clonal CHO cells. Most CHO cells are assessed on their ability to produce the protein of interest over time, known as their stability. But these CHO cells have primarily been studied in bulk, working under the assumption that these bulk samples are identical because of genetic clonality across the sample; however, this does not address other forms of cellular heterogeneity in these ostensibly clonal cells. It is possible these variable stability phenotypes reflect heterogeneity within the clonal samples. In this study, we performed single-cell RNA sequencing on two clonal CHO-K1 cell populations with different stability phenotypes over a 90 day culture period. Our data showed that the instability of the unstable clone was due in part to the emergence of a low-producing subpopulation in the aged samples. This low-producing subpopulation did not exhibit markers of cellular stress which were expressed in the higher-producing populations. Further multiomic investigation should be performed to better characterize this heterogeneity.

## 1. Introduction

To understand the phenotype of cells, we typically rely on measuring RNA levels to give insight into active transcription hence translation. Cells express different genes depending on their age, differentiation status, environmental stimuli, physiological conditions, and even stochastic chance all of which are reflected in a cell’s transcriptome(Kim and Eberwine, 2010). These variations can impact cell behavior and productivity for recombinant cell lines used for industrial biotherapeutic production (Pilbrough et al., 2009). But when performing these methods on a population of cells, measurements can often obscure heterogeneity within the sample; changes within a subpopulation might be missed, or changes in proportions of cell type misconstrued as changes in bulk transcription. The advent of single-cell sequencing, either through split-recombine methods or droplet-based methods, (Cao et al., 2017; Macosko et al., 2015) leverages individual cell oligonucleotide barcodes to allow for computational deconvolution and assignment of each RNA molecule to a cell. With these methods along with computational tools to support them, there has been an explosion in the investigation of heterogeneity within tissues(Cha and Lee, 2020; Choi and Kim, 2019; Deng et al., 2014; Haque et al., 2017; Shalek et al., 2014; Tirosh et al., 2016; Wang et al., 2021).

However such tools are rarely applied to ostensibly “clonal” cell populations. Clonal cell populations, generated from expansion from a single cell, are often assumed to be homogenous, though a cornucopia of data suggests this is not truly the case (Choi and Kim, 2019; Shalek et al., 2014). This is of particular interest in profiling Chinese hamster ovary (CHO) cells, the predominant mammalian expression platform for biotherapeutic production in the biopharmaceutical industry. CHO cells are responsible for producing over 70% of recombinant therapeutic proteins, including monoclonal antibodies, hormones, and vaccines, due to their superior scalability, and well-established regulatory acceptance(Kim et al., 2012). More importantly, their capabilities in post-translational modifications and protein folding are crucial for producing highly compatible human therapeutics (Barnes, 2003).

CHO cells require optimal growth conditions to produce useful amounts of recombinant protein and are prone to genomic instability(Barnes et al., 2003; Dahodwala and Lee, 2019). But in production environments, it is hard to both predict and maintain high growth clones, even with selection pressure. The control mechanisms triggering shifts in response to environmental conditions in these cells are not yet fully understood. Previous transcriptome studies have mainly focused on comparative analyses between different environmental states or defined cell samples at the population level(Clarke et al., 2011; Doolan et al., 2013; Hsu et al., 2017).

The introduction of single-cell RNA sequencing techniques has provided new opportunities to investigate gene expression profiles at the single-cell resolution, offering insights into the sources of variation in cell lines and subclones. However, we argue existing studies have not used enough cells to explore this given the expected proportions of cell populations, or did not study them over culture time. Tzani et al. performed bulk and single cell analysis on 1000 cells in CHO-K1 suspension culture but did not age them to measure changes over time. They did try to reconstruct heterogeneity in their populations with pseudo-temporal ordering, implicating stress-related divergence (Tzani et al., 2021). Ogata et al. (2021) analyzed single-cell transcriptomes of CHO-K1 suspension and adherent cultures, only using ∼100 cells per condition and only aged for <2 weeks (Ogata et al., 2021). Unlike Tzani et al., they observed a correlation between gene expression and cell cycle phase in adherent CHO cells (Ogata et al., 2021). They did not observe other clear substructures, though they did see significant variation in enolase expression within their clusters. Other recent work such as Borsi et al. (2023) compared the transcriptional profiles of differently aged CHO-K1 vs HEK293FT cell lines, specifically looking for highly variable genes shared between the two cell lines (Borsi et al., 2023). Borsi et al. identified 53 genes that defined the transcriptomic variability in both the HEK and CHO cell lines.

But the real concern is not just understanding a snapshot of heterogeneity (Tzani et al., 2021) nor how cells perform over a limited span (Borsi et al., 2023; Ogata et al., 2021) but rather *why production drops over time*. To accomplish this we used 10X Chromium single-cell sequencing to examine the transcriptome of two CHO clones over 90 days (30 population doublings). We explored to see if the transcriptional heterogeneity and subpopulations could act to explain how titer/productivity drops over time.

## 2. Material and Methods

### 2.1. Cell Culture

Two clonal cell lines producing the same monoclonal antibody and their host cell line (CHOZN-GS, SAFC) were provided by Millipore Sigma (St. Louis, MO) as cryo-preserved vials. CHOZN® ZFN-Modified GS-/- platform was used to generate the clones. This CHO strain has the endogenous genes encoding glutamine synthetase(*Glul*) knocked out. The host cells were transfected to introduce a human IgG light chain and heavy chain gene target along with an exogenous glutamine synthetase (*Glul*) gene. Clones were isolated from the pool generated by the above-described method.

These cells were cultured with imMEDIAte ADVANTAGE 87093C (SAFC) media, a serum-free custom made medium by Milipore Sigma, in 125 mL shake flasks (Fisher Scientific) with a working volume of 30mL. Incubation was done in a humidified orbital shaking incubator set at 37C, 80% humidity, 5% CO2 and 125 rpm shaking. The two clonal cell lines were thawed and seeded at 0.3 x 10^6^ cells/mL.

The cells were passaged every 3 days with viable cell density (VCD) and viability was determined by hemocytometer using trypan blue dye exclusion. The cells were then seeded in fresh media at 0.3 x 10^6^ cells/mL. Cells were passaged for a total of 30 passages over 90 days during the aging process. Clones were cultured in triplicate either with or without 6mM L-Glutamine supplementation for the duration of the experiment. Culture samples at passage 0 (P0) (clone A, clone B) and passage 30 (P30) (clone A +Gln, clone A -Gln, clone B +Gln, clone B -Gln) were sampled for 10x cDNA library prep.

Cell banks were established from P30 cultures by freezing 10^7^ cells/mL culture in media supplemented by 10% DMSO. Media exchange was facilitated by pelleting cells at 500xg for 5 minutes and aspirating supernatants. Media exchanged cultures were then aliquoted into cryotubes and frozen in MrFrosty freezing containers (Nalgene) that gradually decreased content temperature in a -80C Freezer. Banked cells were then stored in a -80C freezer for short term storage and in a gas phase LN2 tank for long term storage. To thaw banked cells, we incubated the cryo-vial in a 37C water bath, followed by adding 1 mL of thawed culture to 29 mL fresh media in a shaking flask.

### 2.2. Fed-batch Experiment

Banked clones from the aging experiment were thawed to investigate fed-batch performance. Specifically, we revived clone A at P0 and clone A +Gln/-Gln at P30, as well as clone B at P0 and clone B +Gln/-Gln at P30. They were characterized in triplicate in a 14-day fed-batch process in 125 mL shake flasks (Fisher Scientific). Starting at a seeding density of 0.6 x 10^6^ cells/mL, we cultured them with a working volume of 30 mL with glucose level set at 5.5 g/L. We used Sigma imMEDIAte ADVANTAGE 87093C (SAFC) as media at D0 for all unaged samples as well as samples aged without glutamine(-Glu). The same basal media supplemented by 6mM L-Glutamine was used for samples aged with glutamine(+Glu). Culture conditions were the same as the aging experiment(37C, 5% CO2 and 125 rpm shaking). Glucose was fed into the culture via bolus feed every other day after day 4 of the culture. VCD and viability were recorded daily via hemocytometer. Glucose and lactate measurements were also taken daily using a YSI 2900 Biochemistry Analyzer. Titer was measured from culture supernatant via HPLC run on a POROS Protein-A column. The calibration curve of the HPLC was determined with a manufacturer provided standard.

### 2.3. Library Prep

Approximately 16,000 cells per condition, as measured by hemocytometer, were loaded into individual channels of a Chromium Next GEM Single Cell 3’ v3.1 dual index chip (10X Genomics), each targeting 8,000-10,000 cells. In brief, poly-A mRNA transcripts were captured with gel bead oligo containing poly(dT) sequence, and reverse transcribed into barcoded full-length cDNA. Then, scRNA-seq libraries were prepared with the Chromium Single Cell 3’ library construction kit (10X Genomics), where each sample is amplified with primers containing unique i5 and i7 sample indexes and common P5 and P7 sequencing adaptors via PCR. Libraries were subsequently pooled based on their molar concentration. Six pooled libraries were then loaded at 4nM and sequenced on a NovaSeq SP flow cell (Illumina) with 28 bases for read1, 91 bases for read2, and 10 bases for i5 and i7 index respectively.

### 2.4. Cell Ranger

A Cell Ranger reference was made from the CHOK1GS genome and transcriptome from Ensembl (GCA_900186095.1) using Cell Ranger version 6.0.2. We manually added heavy chain IgG (IgH), light chain IgG(IgL), and the CHO *Glul* gene (NM_001246770) sequences to our reference. We then generated an alignment index from the reference using the *cellranger mkref* command with default settings. This reference was then used with cellranger count, also run using default settings, for all six samples. Barcodes with more than 500 UMIs were considered cells (Supplementary Figure 1).

### 2.5. QC & Filtering

Quality control was done using Scanpy version 1.8.2 (Wolf et al., 2018). Cells with less than 500 genes as well as genes identified in less than 10 cells were removed. We retained cells with mitochondrial gene counts between 0-18%, total transcript counts between 5,000-100,000, and total number of genes between 500-10,000 for subsequent analysis (Supplementary Figure 2). A mitochondrial gene list was obtained from Quiros et al (Quiros et al., 2017). We normalized the remaining cells to 10,000 transcript counts per cell, log1p transformed with base 2, and merged into one single-cell object. Cell cycle regression was then performed using S and G2M phase gene lists to remove cell cycle phase gene variability using the built-in Scanpy regression function (Supplementary Figures 3-4). Human S and G2M gene lists were lifted over onto the K1GS CHO genome (Tirosh et al., 2016).

### 2.6. Highly-variable Gene Detection

Due to the clonal nature of the samples, foreign genes inserted, and limited biological replication. Highly variable genes were selected per sample using the default settings in Scanpy *highly_variable_genes* function. Genes identified as highly variable in 5 or more samples were kept for clustering, resulting in 78 genes (Supplementary Table 1).

### 2.7. Clustering and UMAP

Uniform manifold approximation and projection (UMAP) was calculated using 25 principal components on the selected highly variable genes. Clustering was performed using the Louvain algorithm in Scanpy at a resolution of 0.25. Quality control metrics did not demonstrate any cluster specific bias (Supplementary Figure 5).

### 2.8. Differential Gene Expression

Using the Wilcoxon Rank Sum test, differentially expressed genes were identified between the two IgG expression groups for both clone B P30 samples. Genes with an absolute log2foldchange greater than 1, Bonferroni corrected p-value less than 0.05, average expression greater than 0.2, and non-zero expression percentage greater than 15% were considered significantly differentially expressed. The minimum non-zero expression percentage of 15% was included to remove genes that might appear as differentially expressed due to sparsity of the single cell counts data (Supplementary Figure 6).

## 3. Results

### 3.1. Cell Samples

To characterize changes in cell populations over time, we used two CHO clones provided by our collaborators at Millipore-Sigma, based on the CHOZN® ZFN-Modified GS-/- platform. In these CHO cells the endogenous glutamine synthetase (*Glul*) is knocked out, requiring that their media be supplemented by L-glutamine. These host cells are then transfected to introduce a human IgG with lambda light chain(IgL) and heavy chain(IgH) along with glutamine synthetase (*Glul*). *Glul* allows for selection pressure, as only cells with a successful transfection will be able to grow in the absence of supplemented glutamine. From the pool of transfected cells, individual cells were isolated to form founding clonal populations. We cultured these cells in orbital shaking incubators in a 90 day aging campaign, passing every 3 days for 30 passages either in the presence (no selection) or absence (selection) of 6mM glutamine. Samples from passage 0 and passage 30 were banked in cryogenic storage for subsequent analysis.

We then revived banked cell samples and cultured them in triplicate in a fed-batch system for a maximum of 15 days (Figure 1A). Over the course of the fed-batch culture period, mAb titer, viable cell density (VCD), and cell viability were measured each day (Supplementary Figures 7-9). Plotting titer vs VCD (Figure 1B), we found that samples with high titer typically have a lower VCD and samples with a high VCD a low titer - as might be expected due to the metabolic cost of IgG production. Notably, clone B has a large increase in VCD after aging, independent of selection pressure. At P0, clone A and clone B both showed high titer though clone B was noticeably lower. However, by P30 clone A under selection pressure (-Gln) is the only sample that maintained a high productivity. The other conditions all have significantly reduced titer (<1000 mg/L).

**Figure 1:**
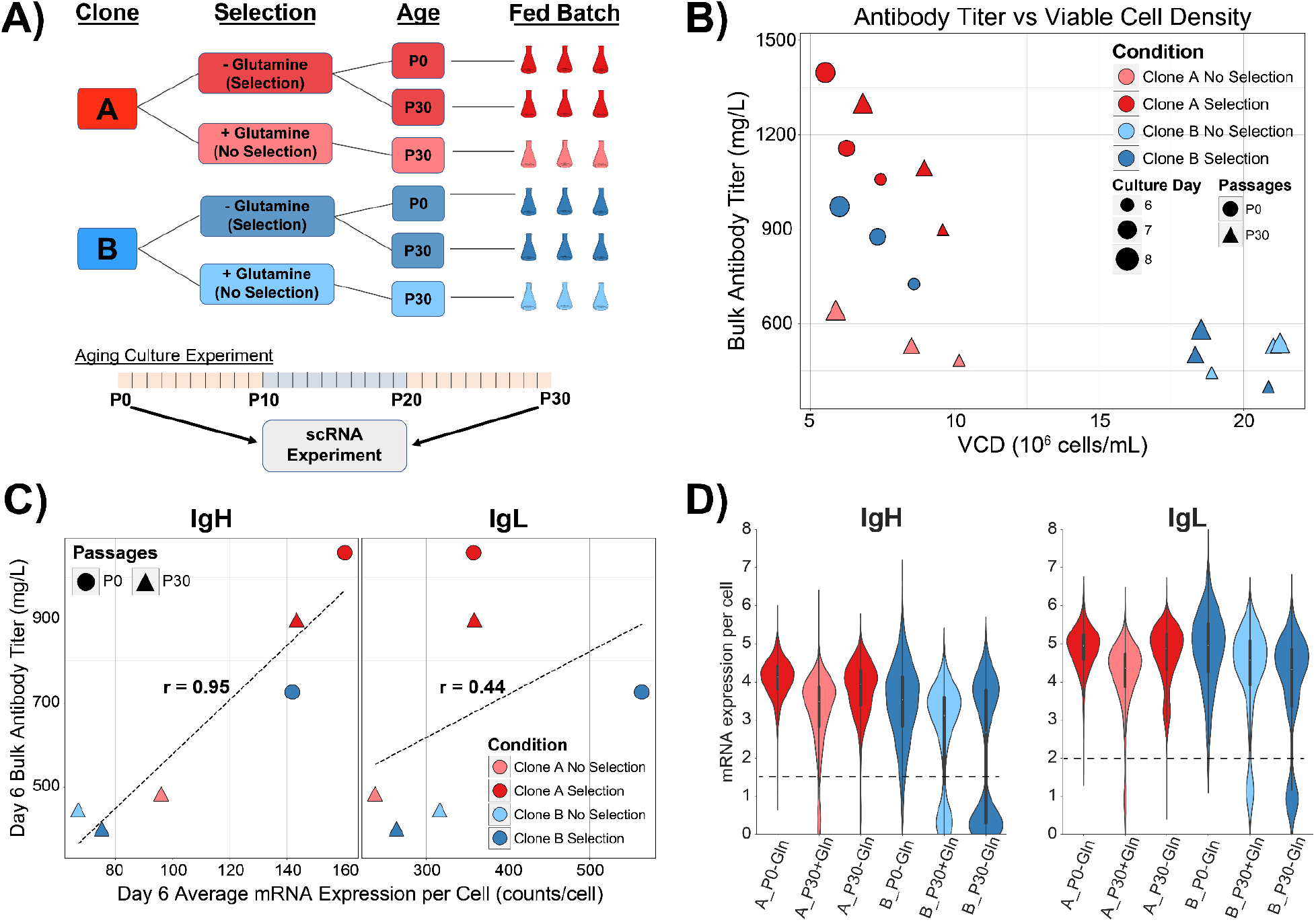
Experimental design. **A)** Two CHO clones with the same gene target were cultured with and without glutamine for 90 days with passages every 3 days. To measure productivity, these six cell samples were cultured in triplicate for measuring mAb titer and cell viability. **B)** Monoclonal antibody titer and viable cell density (VCD) were measured on each day of the cell culture period. Days 6-8 of the fed-batch experiment have VCD plotted vs the bulk titer measurements from the same day. **C)** On day 6 of the fed-batch experiment, the antibody titer and the single cell experiments were performed. For each sample the bulk antibody titer is plotted against the average mRNA transcripts per cell of heavy and light chain IgG. The Pearson correlation coefficient was calculated for both IgG chains. **D)** Violin plots of IgG gene expression per cell per sample. The y-axis is the log2 of the cDNA counts + 1 for each gene per cell. We observe high levels of expression per cell with distinct bimodality in some of the Day 30 samples. Thresholds dividing high vs low IgG expression are indicated by dotted lines

### 3.2. Single cell RNA-sequencing

From day 6 of the fed-batch samples we generated 10X Single Cell Gene Expression profiling data. For each condition, we targeted 10,000 cells in the 10X Chromium controller to generate barcoded cDNA. The cDNA was prepared for Illumina sequencing following established protocols (Methods) and sequenced on a NovaSeq generating an average of ∼636 million reads per sample. Data generated is summarized in Supplementary Table 2. This data was then used to generate a count matrix using 10X software as summarized in Methods.

From this data, we first wanted to measure the levels of target gene expression. We plotted the measured aggregated RNA-sequencing reads for IgG, both heavy chain (IgH) and light chain (IgL) versus the measured titer at day 6 (Figure 1C). We found these values had correlations of R=0.95, and 0.44 for the heavy and light chains, respectively. The high correlation for IgH reflects the nature of the titer assay - it measures IgH levels.

Moving from the bulk RNA expression, we examined the single cell log-transformed expression of the three inserted genes (*IgH, IgL, Glul*) (Figure 1D, Supplemental Figure 10A). We observe a clear bimodality in clone B in both P30 conditions, with a *slight* bimodality of clone A at P30 without selection pressure. Setting a threshold of 1.5 for IgH and 2 for IgL log-transformed expression, we split cells into “high” or “low” expressors. To be a high expressor cells have to pass both thresholds, otherwise, they are classified as low. Notably, clone B P30 samples had 25.7% in P30+Gln and 47.4% in P30-Gln samples exhibiting a low-expressing phenotype (Supplementary Table 3). This suggests that the observed lower IgG protein titer, echoed by the lower bulk RNA level in the B clone is partially due to a subpopulation of cells with lower expression, explaining the variance in viability within the low-titer samples. Curiously, we did not observe the same variation in Glul transcript expression(Supplemental Figure 10), with the caveat that its low levels of expression make it difficult to conclusively rule out variation. The apparent bimodality in the violin plot for Glul is due to a failure to detect *Glul* transcripts due to low expression.

After cell cycle regression, highly variable gene selection, and dimensionality reduction (Methods), we performed Louvain clustering (Figure 2A). Three distinct clusters emerged from the data, each with a unique IgG transcript expression profile (Figure 2B-C). This clustering captures the subpopulations observed in the IgG chain transcript violin plots. Selection pressure acts to maintain a high proportion of productive cells in clone A but seems to have the inverse effect on clone B (Figure 2D), though we observe no difference in Glul expression between the clusters (Supplementary Figure 10B-C). Furthermore, the IgH and IgL expression are not correlated in Cluster 1 which has low IgG heavy chain but high IgG light chain expression. We focused on clone B moving forward due to its high bimodality and presence in the different subpopulations. Differentially expressed genes between Louvain clusters can be found in Supplementary Figure 12.

**Figure 2:**
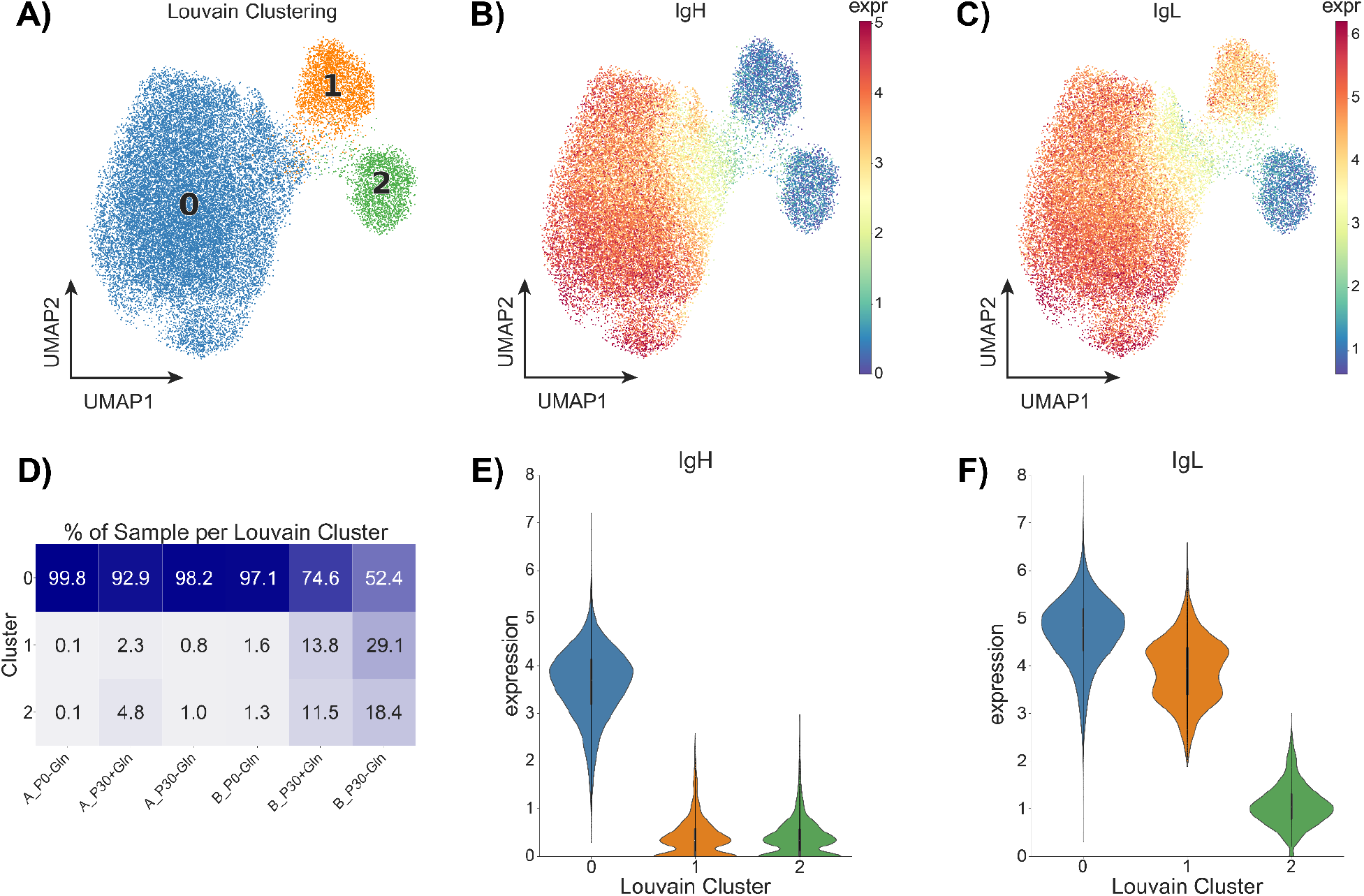
UMAP projections and Louvain clustering of cells from all six samples. **A)** UMAP of cells merged from all six samples colored by Louvain clustering. **B)** UMAP with cells colored by log-transformed IgG heavy chain mRNA expression. Expression is higher in the larger cluster 0 and lower in the two smaller clusters 1 & 2. **C)** UMAP colored by log-transformed IgG light chain mRNA expression; highest in the large cluster, middling in one of the smaller clusters, and low in the other small cluster. **D)** Percent cells from each sample in each Louvain cluster. The clone B P30 samples had a higher presence in clusters 1 & 2, the clusters associated with lower IgG chain expression. UMAP of this data colored by sample can be seen in Supplementary Figure 11. **E-F)** Log-transformed mRNA expression violin plots of the IgG genes by Louvain cluster. Cluster 0 has high IgG heavy chain, and IgG light chain expression. Cluster 1 has low IgG heavy chain and middling IgG light chain. Cluster 2 has a low IgG heavy chain and low IgG light chain.

### 3.3. Digging into Cluster Analysis

To look for biomarkers different between these clusters beyond the target genes, we measured differential gene expression using the Wilcoxon Rank-Sum test between the high IgG producing cluster 0 versus the low IgG producing clusters 1 and 2. We restricted our analysis to clone B P30 samples to remove other clone or age specific artifacts from our analysis. In order to identify truly differentially expressed genes versus noise, we performed multiple filtering steps. Genes where more than 5% of cells had zero expression were removed from differential gene expression analysis; these often have false inflation of either the p-value or log2foldchange when there is actually a marginal difference in non-zero expression between groups(Jiang et al., 2022). Genes were then further filtered to remove genes with average expression < 0.20 (Figure 3B). Finally, genes with magnitude log2foldchange > 1, p-value <0.05, and non-zero expression > 15% were considered as significantly differentially expressed genes (Figure 3A).

**Figure 3:**
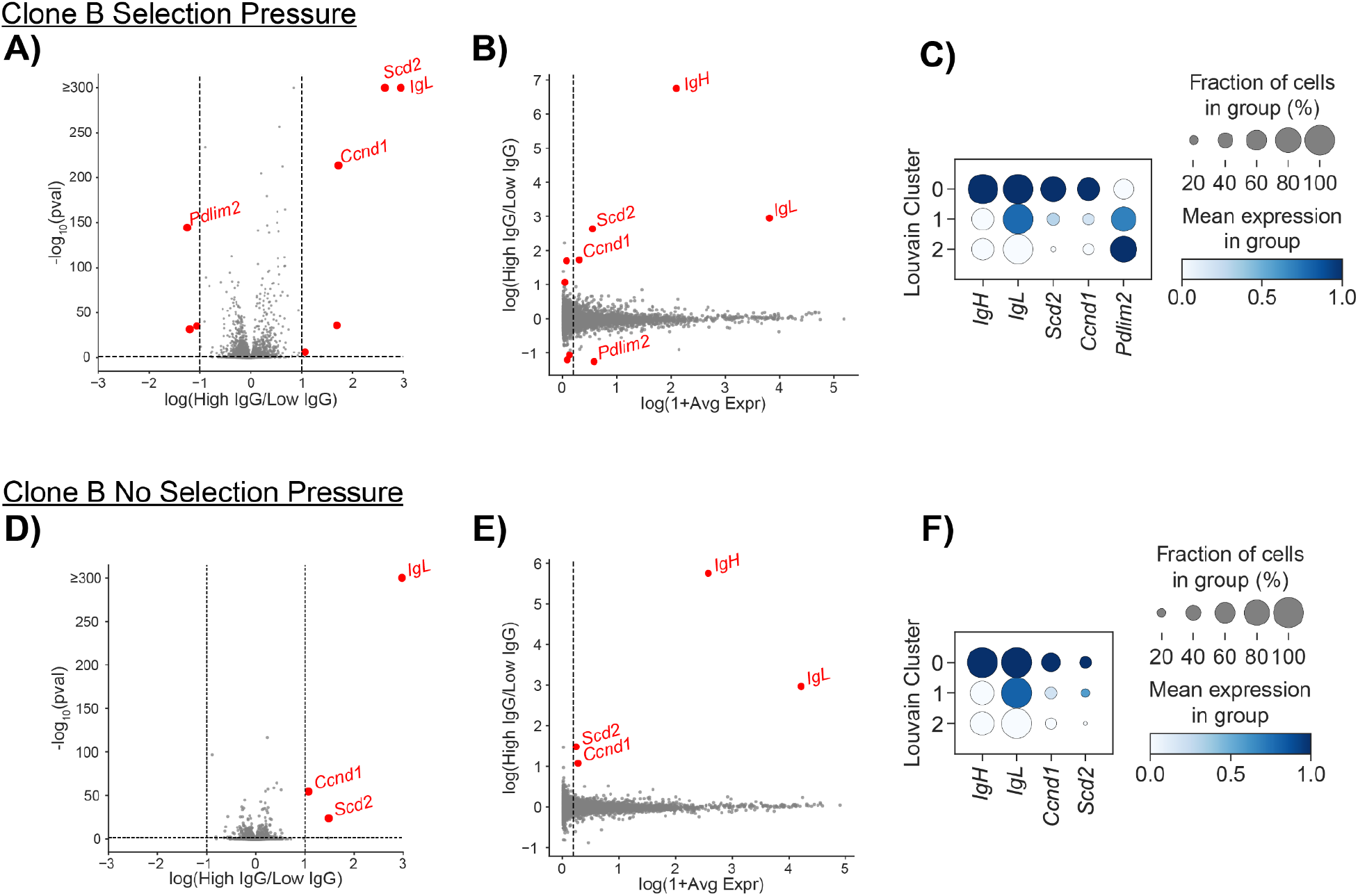
Differentially Expressed Genes between IgG Expression Groups in aged clone B samples. **A)** Volcano plot of genes with p-value generated by Wilcoxon test between the High IgG (Cluster 0) and Low IgG (Clusters 1 and 2) for the clone B P30-Gln sample. Genes expressed in less than 15% of cells were not plotted. Genes with magnitude log2foldchange > 1, p-value <0.05, and non-zero expression > 15% are highlighted in red as significantly differentially expressed genes. Log fold changes are represented as High IgG minus Low IgG. Log p-values were capped at 300 for visualization. X-axis scaling was set from -3 to 3, removing IgH from the plot; full range figure available in Supplementary Figure 13. **B)** Magnitude-Amplitude (MA) plot for all genes in the clone B P30-Gln sample. Genes with average expression greater than 0.2 are labeled in red. The log2foldchange is calculated between IgG expression groups and the average expression is calculated using all data. The five genes that passed all filtering are labeled with red text. **C)** Dot plots of the five significant differentially expressed genes between IgG groups displayed across clusters in the clone B P30-Gln sample. The expression is normalized per gene; size of each dot represents the percentage of cells with non-zero expression for each gene in each cluster. **D)** As **(A)** but for the clone B P30+Gln sample. The same filtering criteria in A) were used. **E)** As **(B)** but clone B P30+Gln sample. **F)** As **(C)** but clone B P30-Gln sample.

This resulted in a set of 5 genes: *IgH, IgL, Scd2, Ccnd1*, and *Pdlim2*. As shown in Figure 3C and 3F, we found that *Scd2* and *Ccnd1* were both highly expressed in Cluster 0 (high production cluster) but lowly expressed or absent from Clusters 1 and 2 (low production cluster). Violin plots of these genes in the clone B P30 samples can be found in Supplementary Figure 14.

We then compared these two genes across all six samples (Supplementary Figure 15). We found that *Ccnd1* was highly expressed in the clone B P0 sample alone, with little difference across the three Louvain clusters. However, *Scd2* was more highly expressed in the four conditions with higher titer and no bimodality in IgG expression (clone A samples, clone B P0). Given this trend, it follows that *Scd2* could be used as a possible biomarker in filtering out cells with low IgG production.

We wanted to see if these differentially expressed genes could act as biomarkers to sort out high versus low expression populations. Through manual inspection we identified a minimum *Scd2* expression threshold of 1.19 transcripts per 10,000 (0.25 in log2 space) for both of the clone B P30 samples. We then filtered out cells with *Scd2* expression below this threshold. We plotted the distributions of *IgH* and *IgL* in Figure 4 before (grey) and after (purple) cell filtering. In the clone B P30-Gln sample, roughly half of the cells (6539 down to 3294, 50.4%) are retained after filtering, but this results in a noticeable increase in average log2 mRNA expression from 2.09 to 3.04 for IgH and 3.81 to 4.51 for IgL. By comparison, the clone B P30+Gln sample retains less cells (7145 to 1708, 24.0%) in filtering and has a less noticeable change in average expression, including no change for IgL and 2.58 to 2.89 for IgH. This filtering strategy works best on the samples under selection pressure even though Scd2 was identified as a differentially expressed gene for both clone B P30 samples. To further investigate this strategy we also applied the filtering to the clone B P0 sample (Supplementary Figure 16). We found that using this *Scd2* threshold most cells were retained (4965 to 4939, 99.5%) and there was minimal change in average expression.

**Figure 4:**
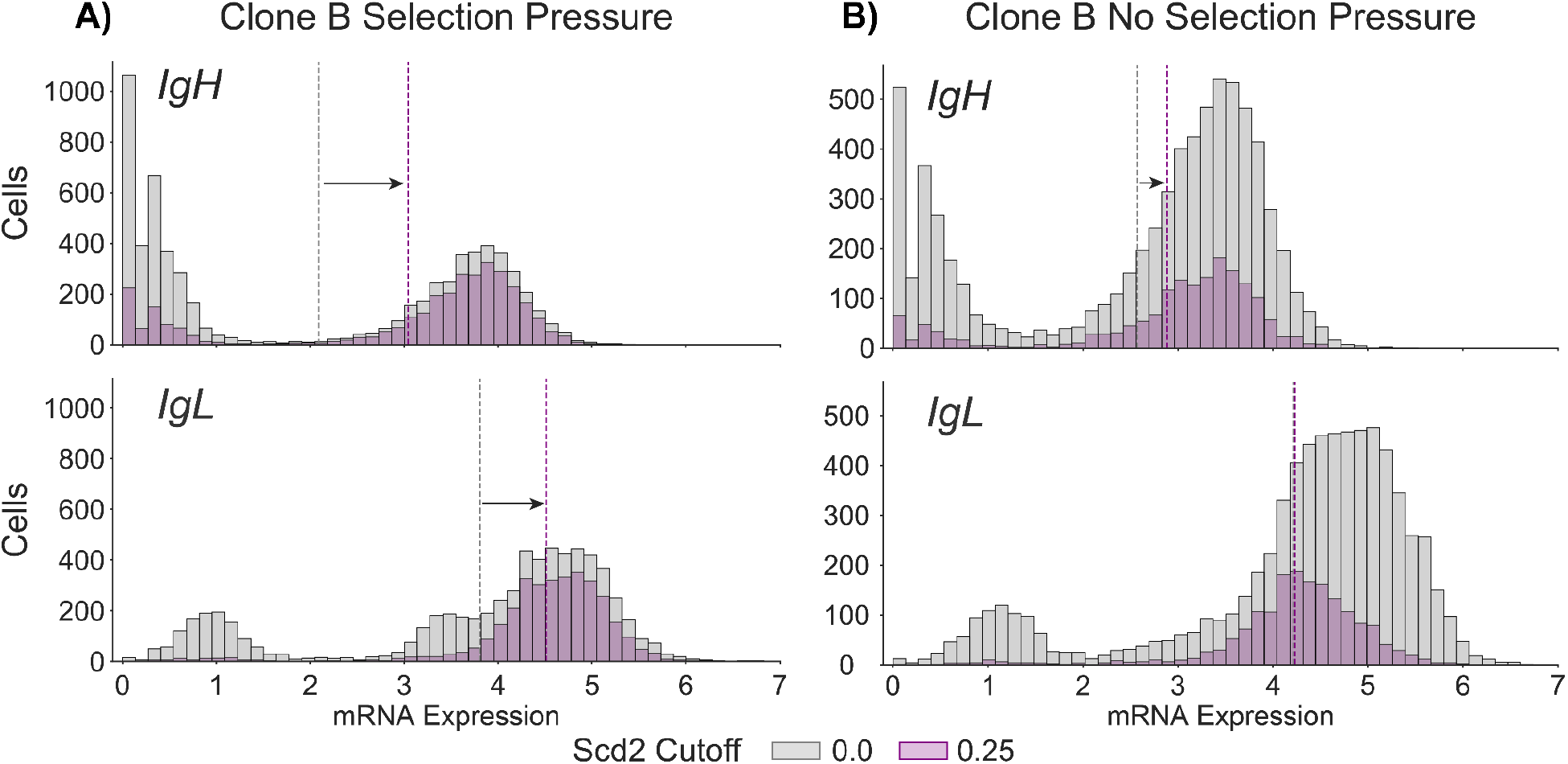
*In silico* cell sorting by coexpressed biomarker *Scd2* in clone B P30 samples. **A)** mRNA expression distributions (grey) of IgH and IgL per cell (n = 6539) are plotted for the clone B P30 sample under selection pressure. Cells were then filtered for a minimum *Scd2* log2 expression threshold of 0.25, with n = 3294 cells passing. IgG expression was then plotted only for these filtered cells (purple). **B)** As in **(A)** but for clone B P30 without selection pressure (n = 7145) and filtered by Scd2 (n = 1708 cells passed threshold). Average mRNA expression per filter condition is displayed as colored vertical dotted lines.

## 4. Discussion

Through our survey of single-cell RNA expression data of two clonal IgG-transfected CHO-K1 cell lines, we could probe the population’s heterogeneity and how it evolved over time correlated with a drop in productivity. We identified three clear subpopulations in our clonal samples using Louvain clustering (Figure 2A). These low producing subpopulations grew as the cells aged, as they are noticeably more populous in the later P30 samples compared to the P0 sample: as high as 47.6% in clone B P30 without Gln, compared to only 2.9% in P0. The two “low expressing” subpopulations differ in their IgG expression phenotype - one has IgG light chain expression but little to no IgG heavy chain expression while the other has neither IgG heavy chain nor light chain expression.

Furthermore, the low IgG expressing cells exhibited a higher cell viability than cells with high IgG expression (Supplementary Figure 8). This suggests that there is an inverse relationship between the cell viability and inserted IgG production. This finding is consistent with the titer and VCD measurements taken from the fed-batch experiment of these same cell lines, as the three samples with highest titer had lower and more consistent VCD than the three samples with lower titer across replicates. It also follows that these three low titer samples had a larger fraction of the low IgG expressing cell subpopulations.

This gain in cell viability at the expense of IgG production leads to the question of how cells in the low-producing subpopulation avoid the selection pressure to better survive. Perhaps most surprisingly, the B P30 sample under selection pressure (B P30-Gln) had a larger portion of low-IgG-producing cells when compared to the B P30 sample without selection pressure (B P30+Gln). In the B P30-Gln sample, the selection pressure seems to have the opposite effect of what one would expect – a lower IgG-producing population than the sample without selection pressure. While we were unable to determine the exact cause of these phenomena in this study, further investigation into this could further explain the gradual reduction in IgG production. We then used the generated single-cell dataset to investigate this question with RNA expression profiles between high and low IgG expression groups across samples.

To understand the differences between these subpopulations, we looked only at the clone B P30 samples which had a large percentage of low-producing cells (Table C3) (25.4%; P30+Gln and 47.6%; P30-Gln). We confined our analysis to the high producing (cluster 0) and low producing (cluster 1 and 2) partitions. We identified differentially expressed genes between the IgG production subpopulations for both of these samples using strict criteria including a p-value cutoff, an absolute log2foldchange minimum, and a minimum percent non-zero expression. The p-value and log2foldchange are standard criteria in finding differential genes; however, the non-zero expression minimum was also crucial in filtering out noisy genes only expressed in a fraction (<10%) of the cells. From our differential gene expression testing, we identified four genes shared between both clone B P30 samples and one gene unique to the B P30-Gln sample. Two of the four shared genes are the IgG heavy and light chain transcripts, a result that reinforces the notion that we divided the cells into IgG production groups properly.

The acyl-CoA desaturase 2 (*Scd2*) and cyclin D1 (*Ccnd1*) genes were similarly coexpressed with the two IgG chains between the production subpopulations. *Ccnd1* is involved in the cell cycle and can be found in either the nucleus or the cytoplasm, functioning differently depending on its location such as DNA damage response and cell cycle regulation (Tchakarska and Sola, 2020). Furthermore, cytoplasmic cyclin D1 has been indicated in metabolic regulation of cells, possibly relating to the successful adaptation to the GS system (Tchakarska and Sola, 2020). We do not believe this is an artifact of cell cycle, as we regressed out cell cycle, and subsequent analysis does not show any cluster-specific cell cycle principle components (Supplemental Figures 3-4). *Scd2* is located in the endoplasmic reticulum and is involved in fatty acid synthesis pathways (Zhou et al., 2021). *Scd* overexpression has been shown to have a role in plasma membrane content and therefore alter the membrane flexibility (Sun et al., 2003). It also plays a key role in mitochondrial metabolism and mTOR activity which would typically improve protein production. These two genes may serve as a cellular readout of stress accompanying the protein production and, when under selection pressure, glutamine synthesis. Using these differentially expressed genes as biomarkers, we tried an *in silico* cell sorting experiment to determine if we could select for the high producing subpopulation by looking for cells with high levels of *Scd2*. We found that we could effectively filter low-expressing cells out of the population using *Scd2* levels but primarily in the aged B P30 samples. Nonetheless, this criterion could be valuable in obtaining subclones with longer sustained production in large scale cell culture bioprocesses. Furthermore, this approach seems to work better when the cells are under selection pressure.

## 5. Conclusions

Our findings reinforce the notion that high-IgG-producing cells are under higher stress than the low-producing subpopulation. Given the higher survivability, lower IgG titer, and lack of defining stress response genes, the low producing subpopulations seem to have subverted the selection pressure and are trending toward cellular stability. We believe that the protein product of genes, in our case *Scd2*, which identifies the low-expression population can be used as cellular markers in cell sorting or other forms of continued selection. Despite these findings, further investigation should be performed to validate and ultimately applythe differential genes and biomarkers identified.

## Acknowledgments

Thanks to all AMBIC member universities (Johns Hopkins University, Clemson University, University of Delaware, University of Maryland College Park, and the University of Massachusetts Lowell) and companies for their mentorship and financial support. Thanks to MilliporeSigma for sharing CHO cells. This work was supported by IUCRC NSF Grant 1624684.

