## Supplemental Figures for "Investigating Subpopulation Dynamics in Clonal CHO-K1 Cells with Single-Cell RNA Sequencing"

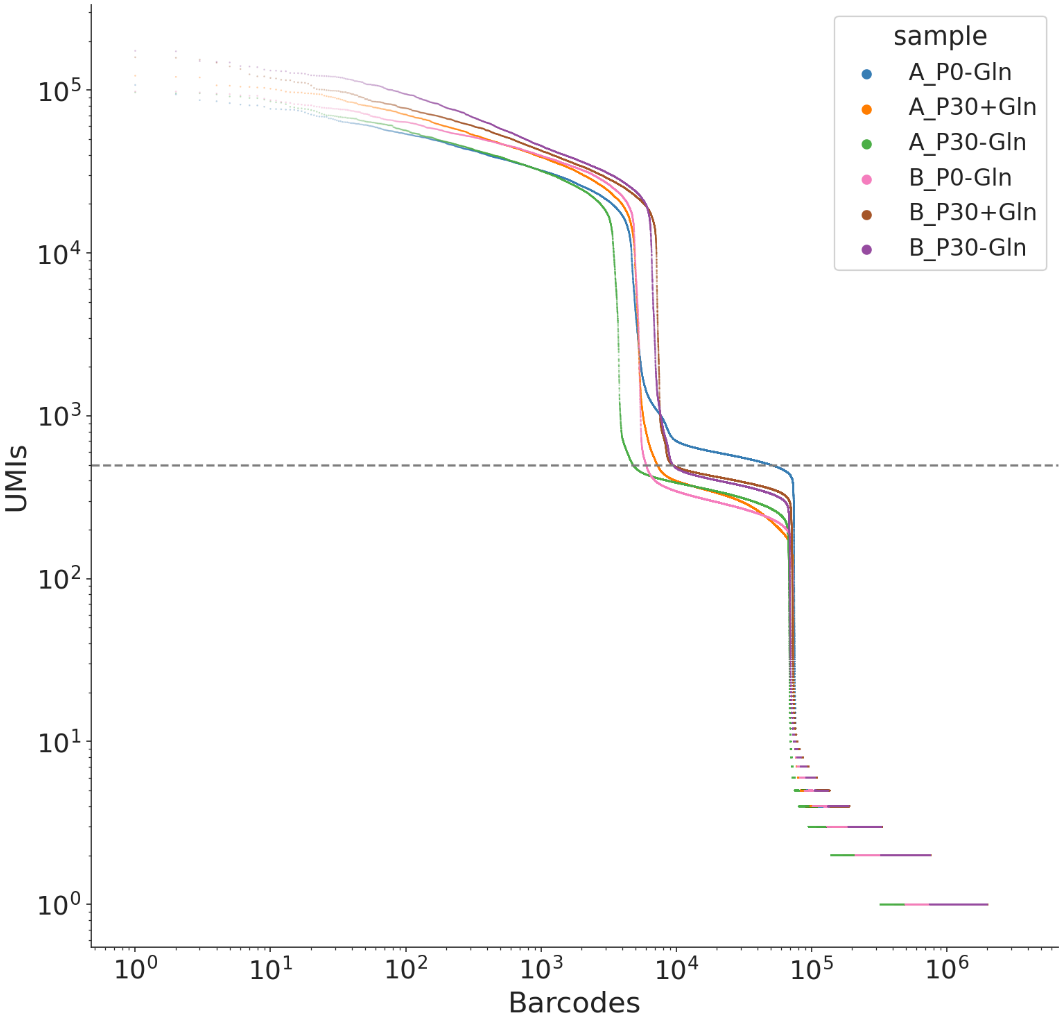


**Supplementary Figure 1: Knee plot of single cell data.** The knee plot of the UMIs vs barcodes is plotted for the six samples of the Fed-Batch scRNA sequencing experiment. Cell barcodes with more than 500 UMIs used for subsequent analysis.

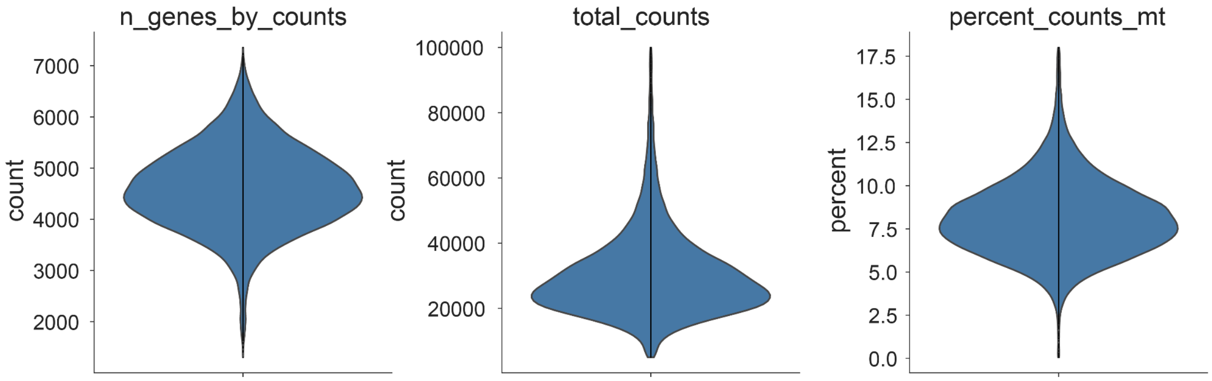


**Supplementary Figure 2: Single cell quality control metrics post filtering.** The Scanpy standard quality control metrics across all samples. *N_genes_by_counts* represents the number of unique genes present per cell after UMI collapsing. The *total_counts* represents the total total number of unique reads per cell. Finally, *percent_counts_mt* represents percentage of reads per cell mapping to identified mitochondrial genes.

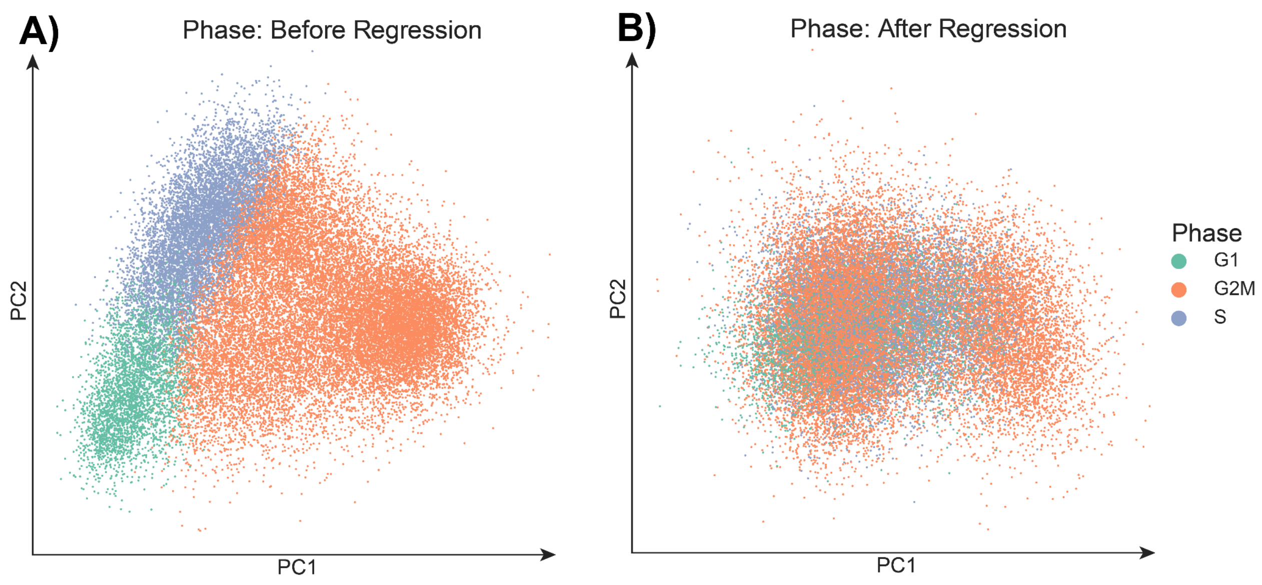


**Supplementary Figure 3: Cell Cycle Regression PCA plots.** The first two principal components of the cell cycle genes are plotted before and after the default Scanpy cell cycle regression function. Each dot represents a cell and is colored by the highest scoring cell cycle phase assigned in the regression function.

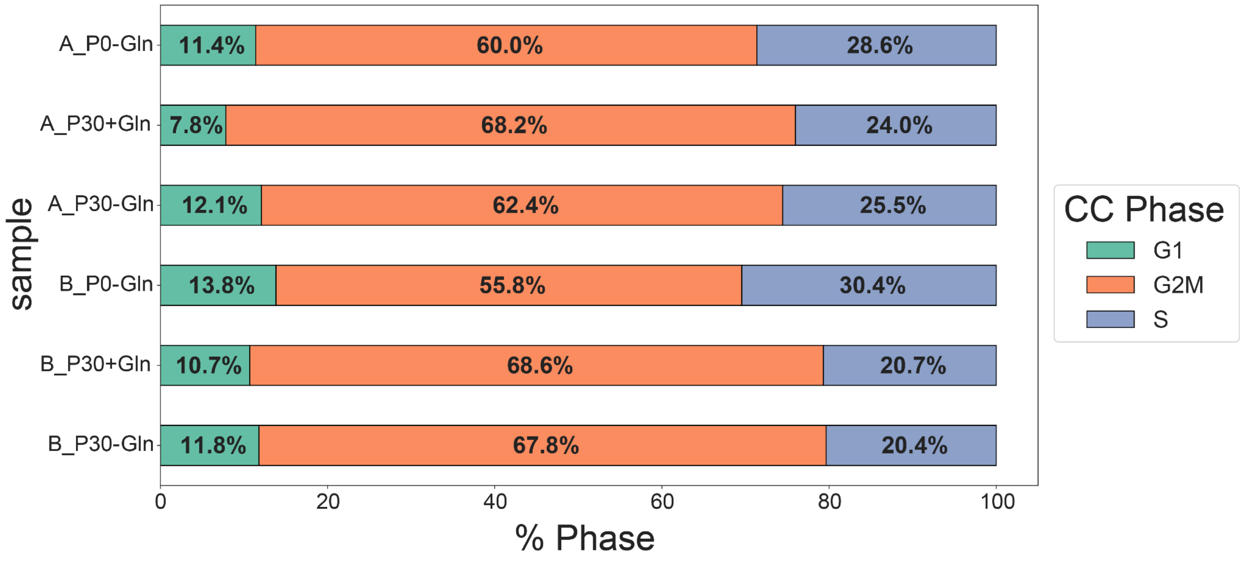


**Supplementary Figure 4: Cell Cycle percentage per sample.** Above are the relative percentages of the cell cycle state for each of the six samples after cell cycle regression. Cells are assigned a cell cycle state in the regression function based on scoring of annotated cell cycle gene lists. The majority of the cells are identified to be in the G2M state. As the cells age, there seems to be a shift to G2M state at the equal expense of the other two cell cycle states.

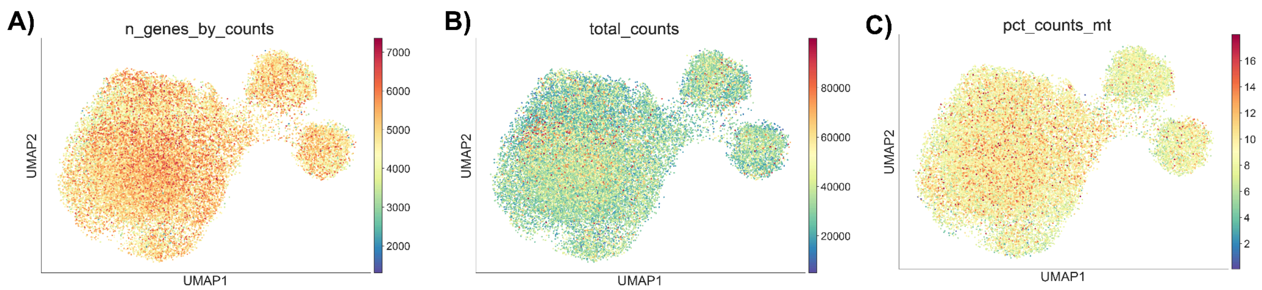


**Supplementary Figure 5: UMAP plots of single cell transcriptomic quality control data. A), B), C)** The three Scanpy single-cell quality score metrics found in Figure S4 are plotted over the UMAP of the highly variable genes.

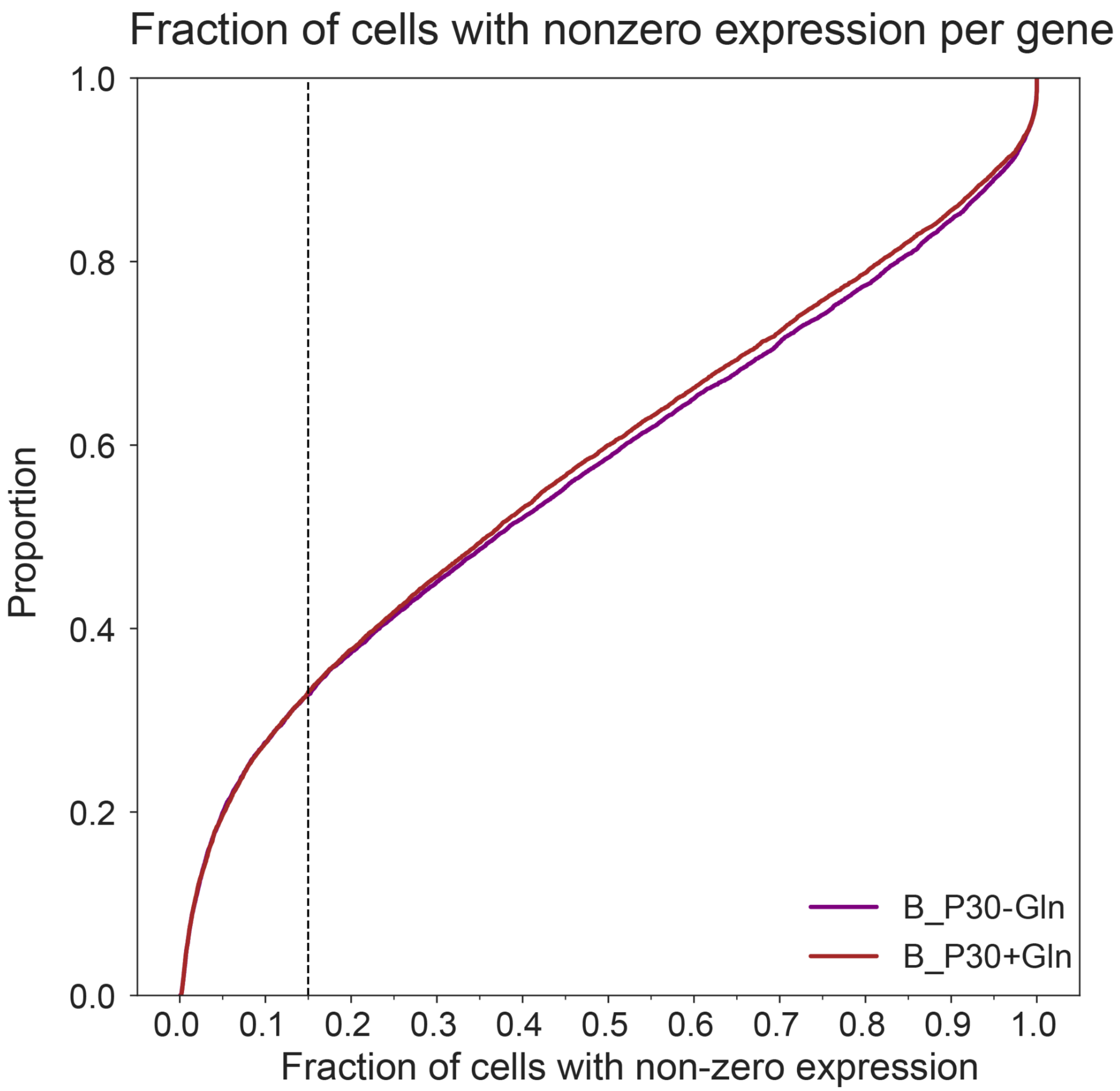


**Supplementary Figure 6: ECDF plot of cells with non-zero expression in the Clone B P30 samples.** The percentage of cells with non-zero expression is calculated per gene per sample. The empirical cumulative density function of these values are plotted per sample in order to identify bias in differential gene expression truly caused by sparsity of the gene. A threshold was set at 0.15 non-zero expression for differential gene expression.

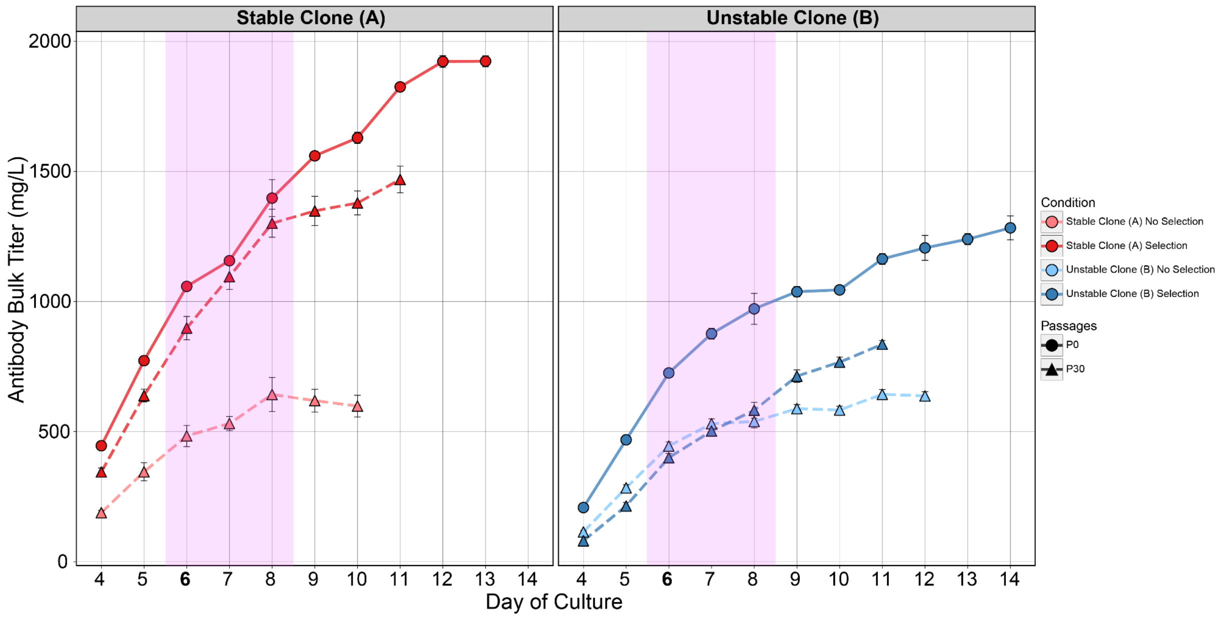


**Supplementary Figure 7: mAb titer of fed batch over culture period.** The bulk antibody titer is plotted over the culture period for the fed-batch experiment. Samples on day 6 used for the single-cell experiment. The pink box highlights days included in Figure 1.

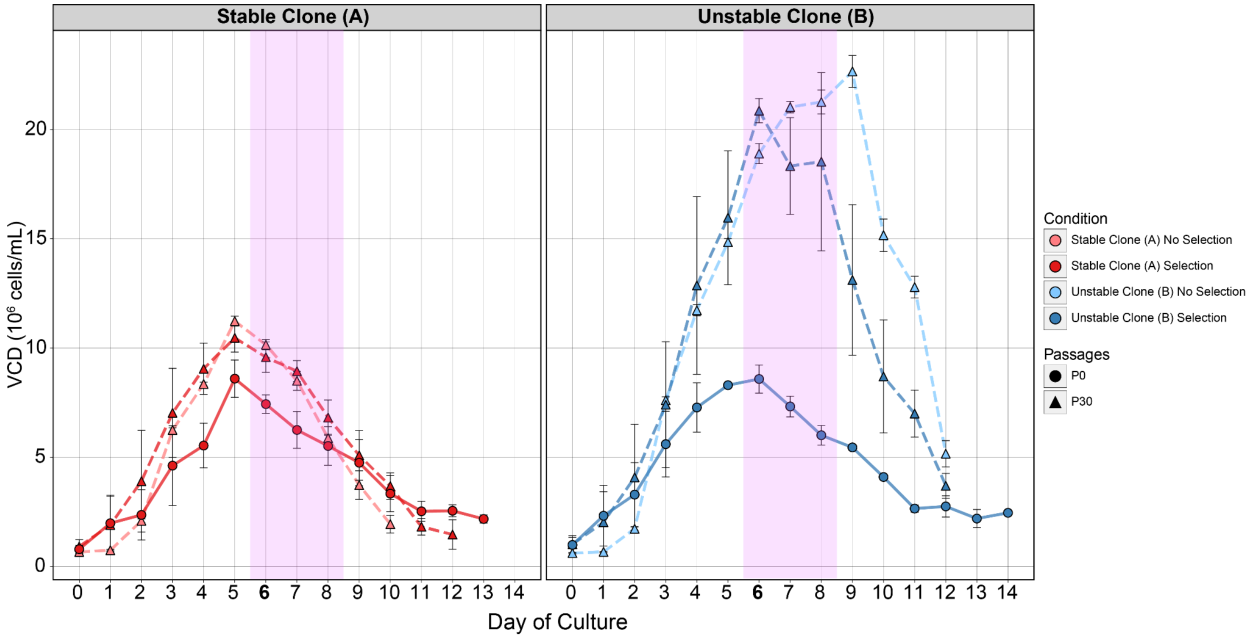


**Supplementary Figure 8: Viable Cell Density (VCD) for fed batch over the culture period.** Viable cell density is plotted over the culture period for the fed-batch experiment. Samples on day 6 used for the single-cell experiment. The pink box represents days included in Figure 1.

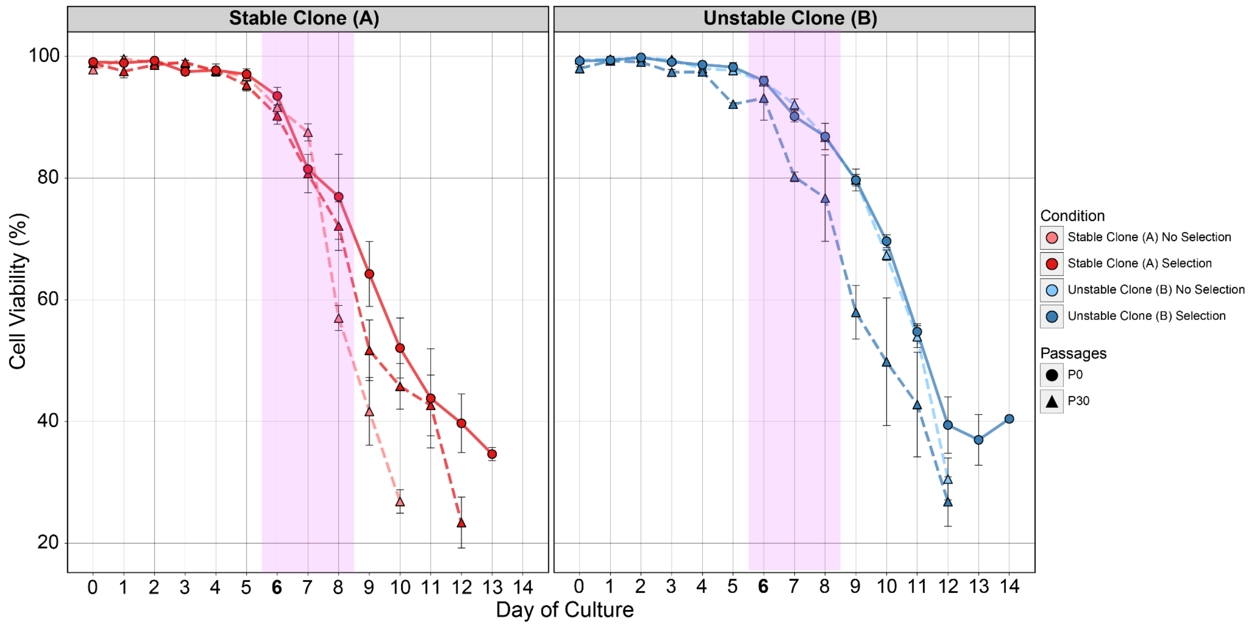


**Supplementary Figure 9: Cell Viability for fed batch over the culture period.** The cell viability is plotted over the culture period for the fed-batch experiment. Samples on day 6 used for the single-cell experiment. The pink box represents days included in Figure 1.

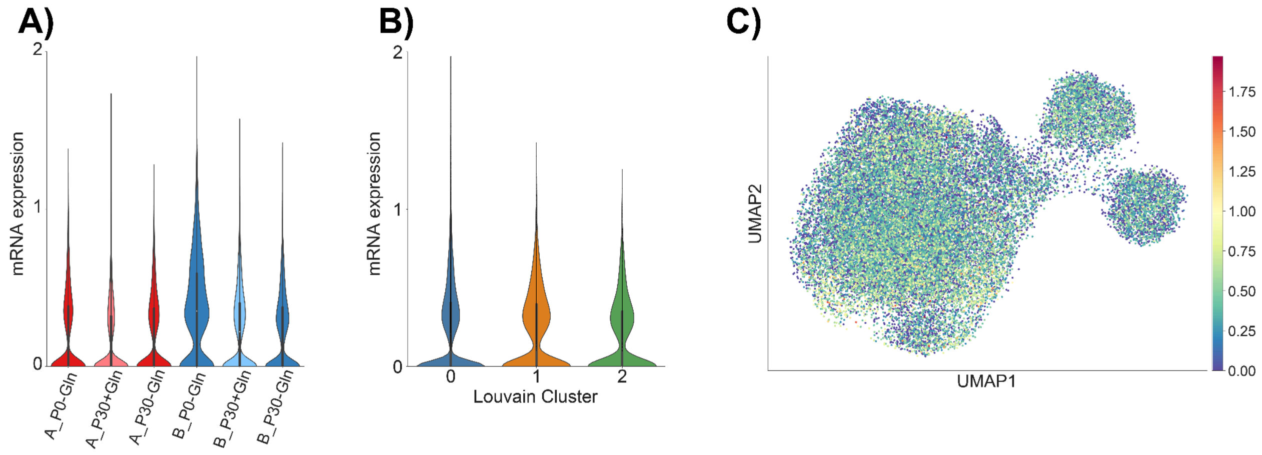


**Supplementary Figure 10: *Glul* mRNA expression profiles.** The per cell Glul transcript expression distribution is plotted **A)** for all six samples, **B)** each Louvain cluster, and **C)** for each cell in a UMAP embedding.

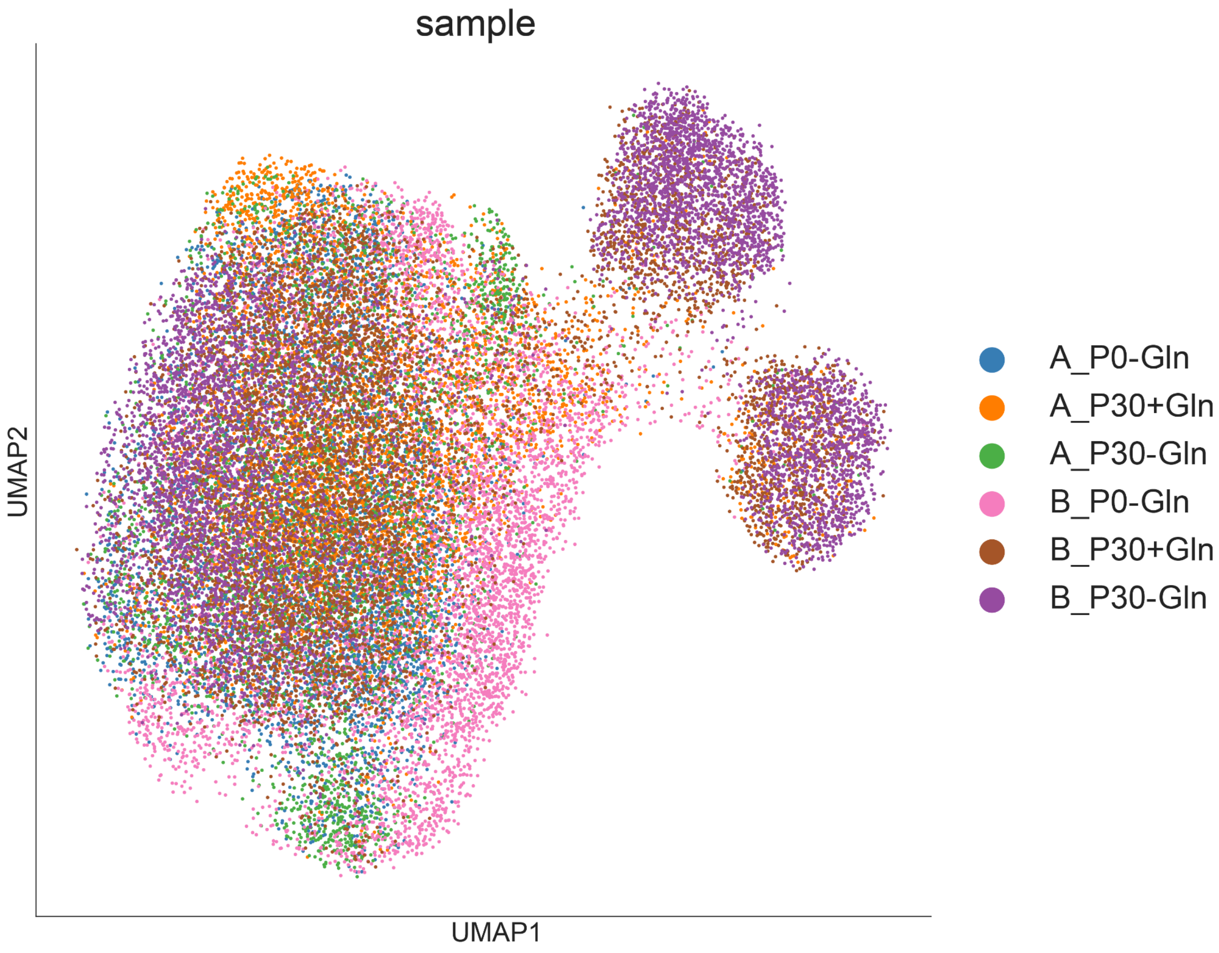


**Supplementary Figure 11: UMAP plot of single cell transcriptomic data per sample.** The sample of origin per cell is plotted on the UMAP visualization.

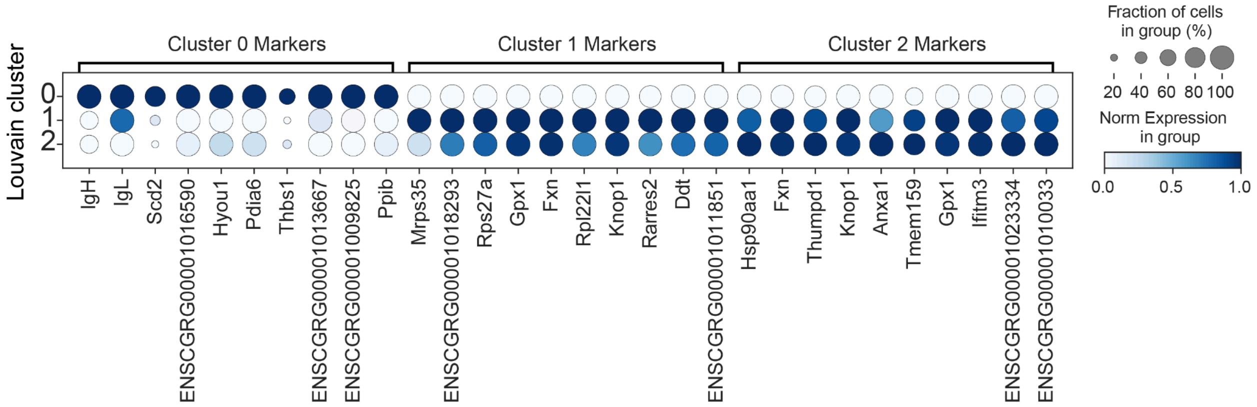


**Supplementary Figure 12: Dotplots of differentially expressed genes between Louvain clusters.** Dot plots of 10 significant differentially expressed genes between Louvain clusters for all six samples. The expression is normalized per gene and size of each dot represents the percentage of cells with non-zero expression for each gene in each cluster.

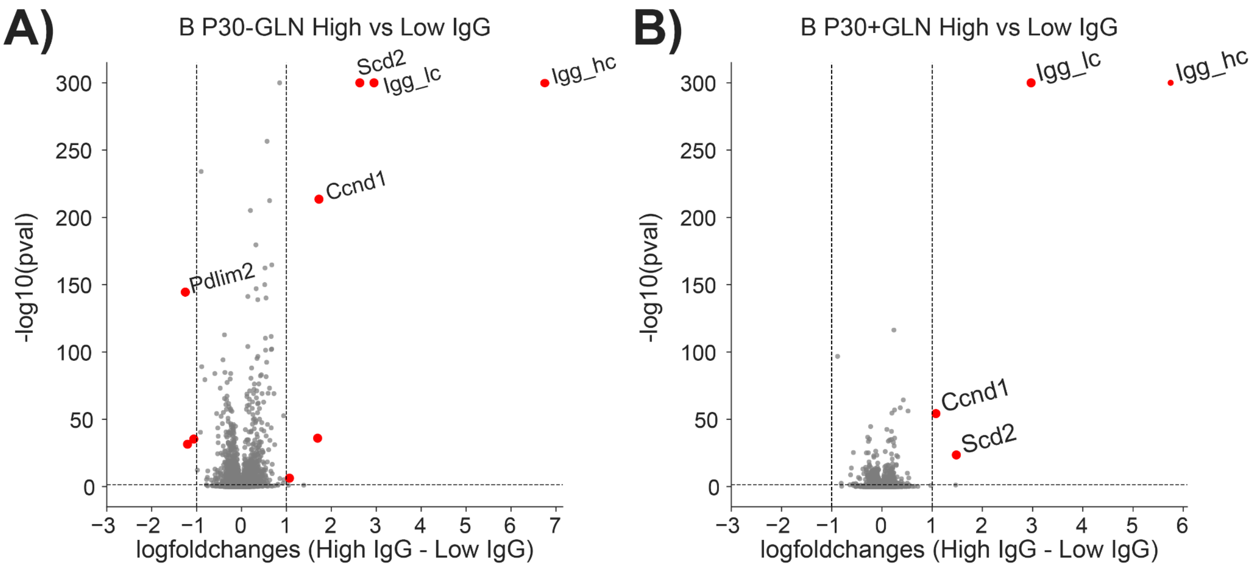


**Supplementary Figure 13: Volcano plots of differentially expressed genes in the Clone B P30 samples.** Differential gene expression was performed between Clusters 0 (high IgG expressing) and Clusters 1+2 (low IgG expressing). The volcano plots of the differential gene expression are plotted for both the Clone B P30 samples. Genes with black dots and red text have passed filtering criteria and considered valid differentially expressed genes.

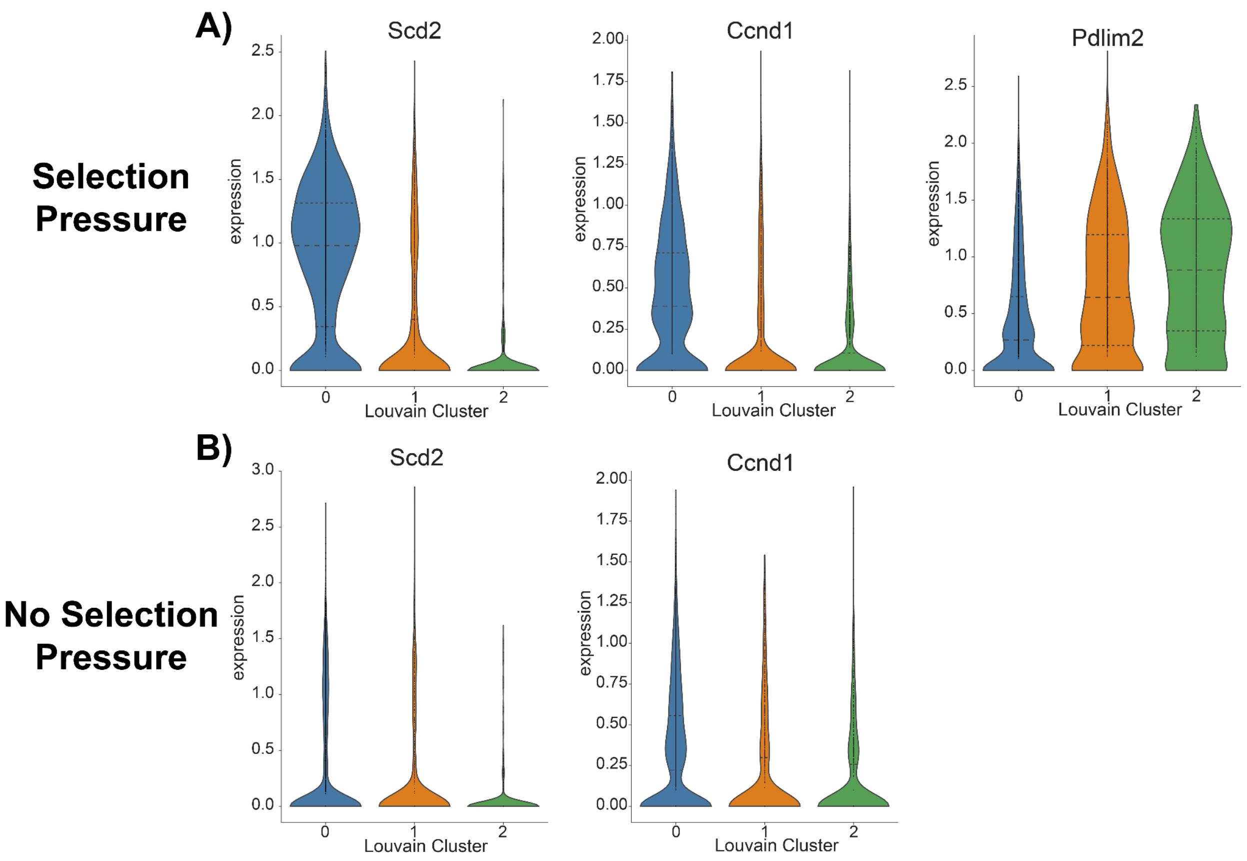


**Supplementary Figure 14: Violin plots of differentially expressed genes for Clone B P30 samples. A)** Violin plots of the log1p-transformed expression of the five significant differentially expressed genes between IgG groups are plotted for the Louvain clusters in the sample. We expect to see the two IgG chains here as that is the defining feature between groups; however, we also see *SCD2* and *CCND1* coexpressed with the IgG chains and *PDLIM2* inversely related with IgG chain expression. **B)** Violin plots of the log1p-transformed expression of the five significant differentially expressed genes between IgG groups are plotted for the three Louvain clusters. We expect to see the two IgG chains here as that is the defining feature between groups; however, we also see *SCD2* and *CCND1* coexpressed with the IgG chains and *PDLIM2* inversely related with IgG chain expression.

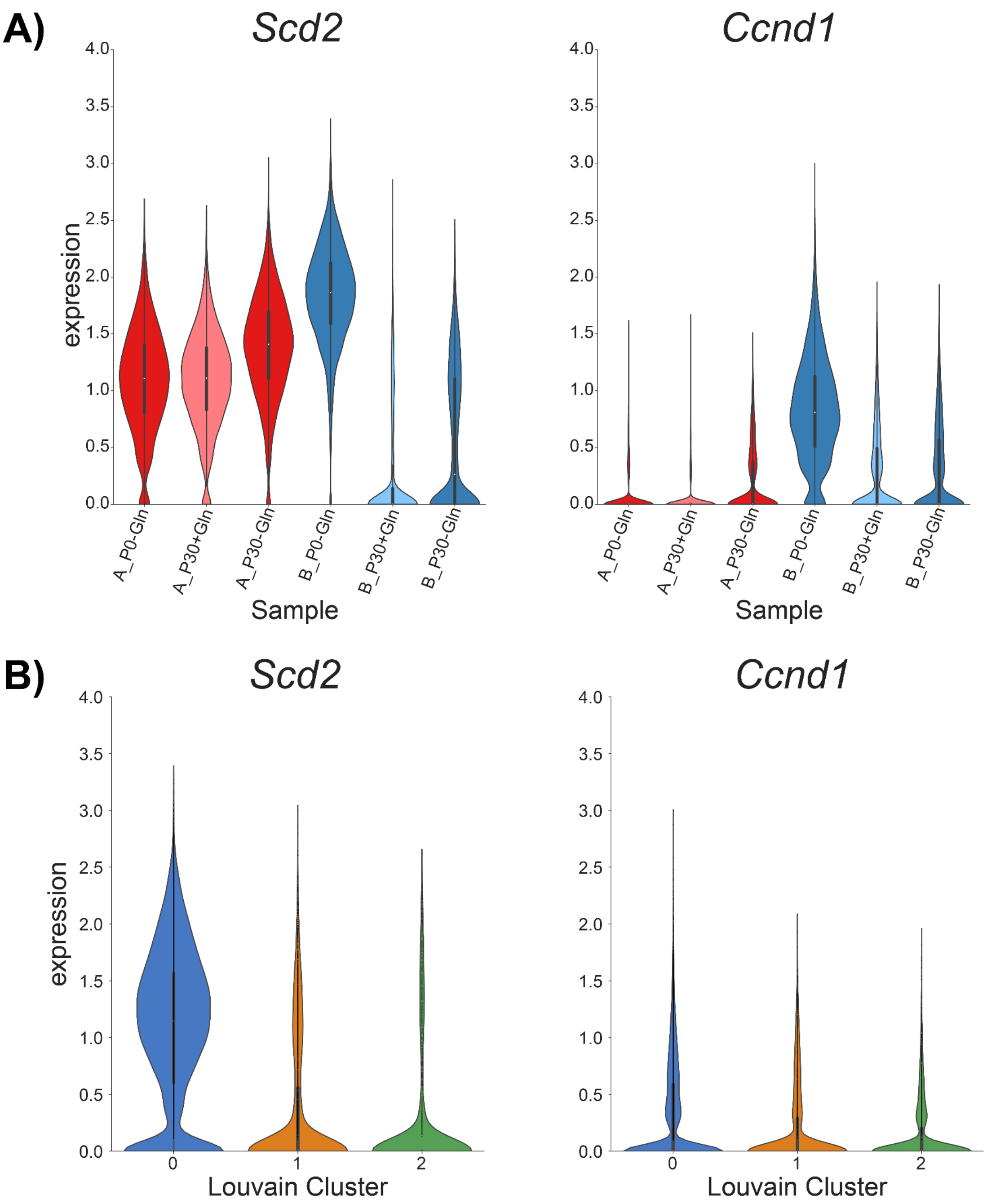


**Supplementary Figure 15: Upregulated genes across all samples. A)** Violin plots of the log1p-transformed expression of the two upregulated genes for all samples are plotted per sample for all six samples. **B)** Violin plots of the log1p-transformed expression of the two upregulated genes for all samples are plotted per Louvain cluster using data from all six samples.

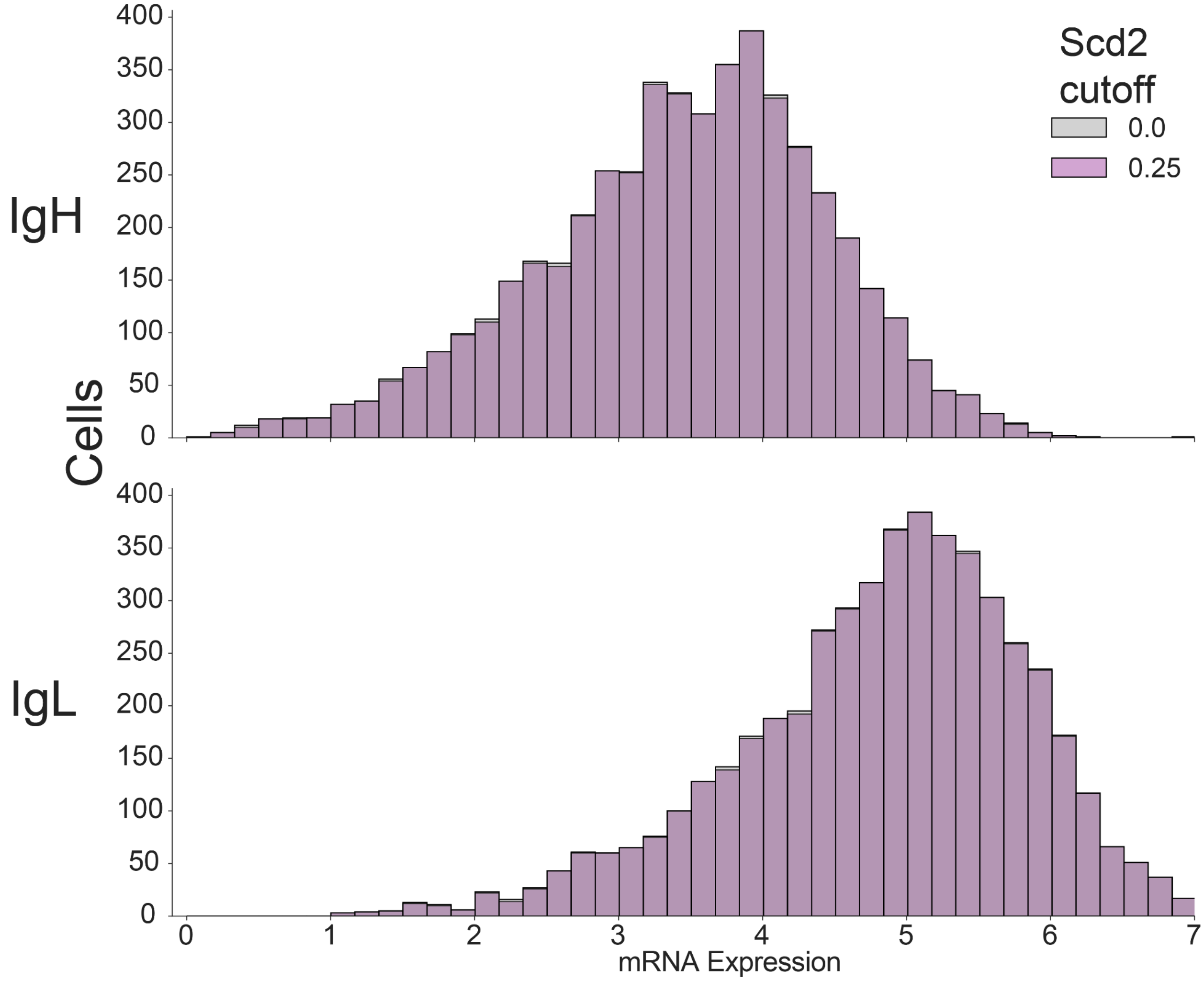


**Supplementary Figure 16: *In silico* Cell sorting by coexpressed biomarker *Scd2* in clone B P0 sample.** mRNA expression distributions (grey) of IgH and IgL per cell (n = 4965) are plotted for the Clone B P0 sample under selection pressure. Cells were then filtered for a minimum *Scd2* log2 expression threshold of 0.25, with n = 4939 cells passing. IgG expression was then plotted only for these filtered cells (purple).
